# Longitudinal cell-free DNA methylome and fragmentome profiles in health uncover signatures of cell type and demographic origin

**DOI:** 10.1101/2025.10.25.684551

**Authors:** Mio Aerden, Tatjana Jatsenko, Kaat Leroy, Kobe De Ridder, Anna Nootens, Valentina Piatti, Koen Devriendt, Joris Robert Vermeesch, Huiwen Che, Bernard Thienpont

## Abstract

Cell-free DNA (cfDNA) is a powerful analyte for liquid biopsy applications. However, the composition and fragmentation of cfDNA in health remains incompletely characterized. Here, we profiled 432 plasma cfDNA samples from healthy individuals using targeted enzymatic methyl-sequencing, allowing cell-type of origin inference and assessment of fragmentation features. Both in a diurnal and a cross-sectional cohort, we observed that cfDNA levels and cellular contributions show lower variability within than between individuals. Hematopoietic lineages are the dominant cfDNA sources, with interindividual variability particularly evident in granulocyte contributions. Demographic factors such as sex, age and body mass index (BMI) contribute to changes in the contribution of various blood cell types, and cfDNA concentrations are 1.6-fold higher in early morning collections (*P* = 1.8 × 10^-5^). By integrating cell-type-specific cfDNA methylation, fragment size and dinucleotide end motif information, we demonstrate distinct associations of these characteristics with specific cell types. Granulocyte-derived fragments showed a consistent enrichment in mononucleosomal sizes (*P* = 3.5 × 10^-54^) and CC end motifs, while also cfDNA from non-hematopoietic cells exhibited distinct size and end-motif profiles. In addition, we observed hypomethylated DNA to be associated with shorter fragment sizes and altered end-motif frequencies, emphasizing interactions between DNA methylation, nuclease activity and chromatin context in shaping cfDNA features. Together, our results provide a view on processes and cell types involved in cfDNA biogenesis in healthy individuals. They underscore that demographic variables and sampling time should be considered for cfDNA-based assay design, but also highlight novel opportunities to improve the representation of specific cell types in cfDNA, thus providing a foundation for optimizing cfDNA diagnostics by leveraging multiple axes of information.

## INTRODUCTION

Cell-free DNA (cfDNA) is released into the bloodstream either as a byproduct of cell death or through active secretion, and is cleared by the liver and spleen, by passive kidney filtration or by direct nuclease-mediated degradation^1,2^. cfDNA has been used as a non-invasive biomarker enabling disease detection, monitoring, and risk assessment in various clinical settings^3,4^. Despite its increasing use in the clinic, the effects of demographic and physiological factors on cfDNA characteristics remain underexplored. A variety of factors, including age, sex and body mass index (BMI) can influence the turnover of cells and the processing of the DNA they release^5,6^. By affecting cfDNA, these demographic variables can introduce interindividual variability, which may mask or confound disease-specific signals and affect the interpretation of cfDNA-based diagnostics. There is hence a need for comprehensive and in-depth analyses of these factors.

Advances in DNA sequencing and analysis have enabled a detailed characterization of cfDNA fragments. cfDNA for example retains the methylation patterns of its cells-of-origin, and methylation signatures found in cfDNA can predict pregnancy complications or detect cancer^7–10^. Additionally, deconvolution analyses of cfDNA methylomes can quantify the proportional contribution of different cellular sources to a heterogeneous cfDNA mixture^11–16^. cfDNA is also fragmented in a specific manner, yielding characteristic patterns shaped by nucleosome positioning and nuclease activity. These cfDNA fragmentation patterns can reflect tissue- and disease-specific differences^17–19^. Fragment size distribution, typically peaking around 167 base pairs (bp) in plasma, can vary across physiological and disease contexts. For instance, in the blood plasma of pregnant women, placenta-derived cfDNA is typically shorter than maternal cfDNA^20^. Beyond fragment size, cfDNA end motifs that are defined by the nucleotides at 5’ termini, provide further insights into the biology of cfDNA degradation: these motifs reflect nuclease cleavage preferences, and their analysis can reveal disease-specific fragmentation signatures^21,22^. Furthermore, while cfDNA features such as methylation and fragmentation are often analysed in isolation, emerging research has demonstrated notable interactions between both^23,24^. However, these modalities are often treated as parallel sources of information, leaving their interdependencies poorly understood and limiting insights into their biological coordination. A systematic investigation of biological heterogeneity and correlations across cfDNA characteristics is hence of particular interest.

The majority of plasma cfDNA originates from different hematopoietic cell types, while contributions from other non-hematopoietic cells have been suggested^11,12,25^. Whether cfDNA from these different cellular sources undergoes differential processing remains an open and largely unexplored question. While cfDNA can be unequivocally assigned to specific tissue types when genetic differences are present, such analyses are only feasible in specific contexts such as cancer, pregnancy or organ transplant where genetically distinct cfDNA populations exist^6^. DNA methylation patterns offer an alternative for inferring the cellular origins of cfDNA. We therefore set out to perform an in-depth analysis of cfDNA obtained from a cohort of healthy individuals, to assess how cfDNA characteristics are affected by demographic and sampling-related factors, as well as by the cellular composition of cfDNA in plasma. Using deep targeted enzymatic methylome sequencing of 432 samples, we characterized variations in cfDNA concentration, methylation and fragmentation patterns across individuals and evaluated the associations between these cfDNA attributes.

## RESULTS

### Diurnal variation in cfDNA composition

First, we wanted to assess whether the sampling moment affects cfDNA characteristics. We recruited 16 healthy individuals between 20 and 30 years old (Table 1) and collected their blood plasma from venipunctures in the morning, afternoon, and evening for three consecutive days, yielding 144 samples (9 per individual; Figure 1a; Supplementary Table 1). From these, we generated sequencing libraries using enzymatic methylome sequencing (EM-seq), a non-destructive method that ensures accurate cfDNA methylation (cfDNAme) profiling while maintaining native fragment sizes and end motifs. We next performed in solution capture with custom probes targeting 4,991 genomic regions of interest (Methods), followed by sequencing to a median on-target depth of 91-fold.

**Table 1:** Overview of diurnal and cross-sectional cohort characteristics. BMI: body mass index. BMI is shown as the Median [Q1;Q3].

| cohort | participant demographics |  |  |  |  | health status |  |
| --- | --- | --- | --- | --- | --- | --- | --- |
|  | participants | female/male | BMI (kg/m<sup>2</sup>) | never smoker | regular high | cancer | prescribed |
| **diurnal** | 16 | 8/8 | 23.11 [21.1;24.3] | 13 (81%) | 9 (56%) | 0 | 5 (31%) |
| **cross-sectional** | 144 | 78/66 | 23.9 [21.8;25.8] | 111 (77%) | 74 (51%) | 11 (8%) | 60 (42%) |
| **20-30** | 21 | 11/10 | 22.1 [20.6;23.8] | 20 (95%) | 16 (76%) | 0 | 7 (33%) |
| **31-40** | 24 | 13/11 | 22.7 [21.1;25.5] | 20 (83%) | 16 (67%) | 1 (4%) | 8 (33%) |
| **41-50** | 25 | 14/11 | 24.3 [20.7;26.9] | 23 (92%) | 12 (48%) | 2 (8%) | 6 (24%) |
| **51-60** | 24 | 13/11 | 24.3 [22.4;26.5] | 19 (79%) | 13 (54%) | 3 (13%) | 10 (42%) |
| **61-70** | 24 | 14/10 | 24.6 [22.7;26.0] | 14 (58%) | 5 (21%) | 3 (13%) | 10 (42%) |
| **>70** | 26 | 13/13 | 25.6 [22.8;26.5] | 15 (58%) | 12 (46%) | 2 (8%) | 19 (73%) |

**Figure 1:**
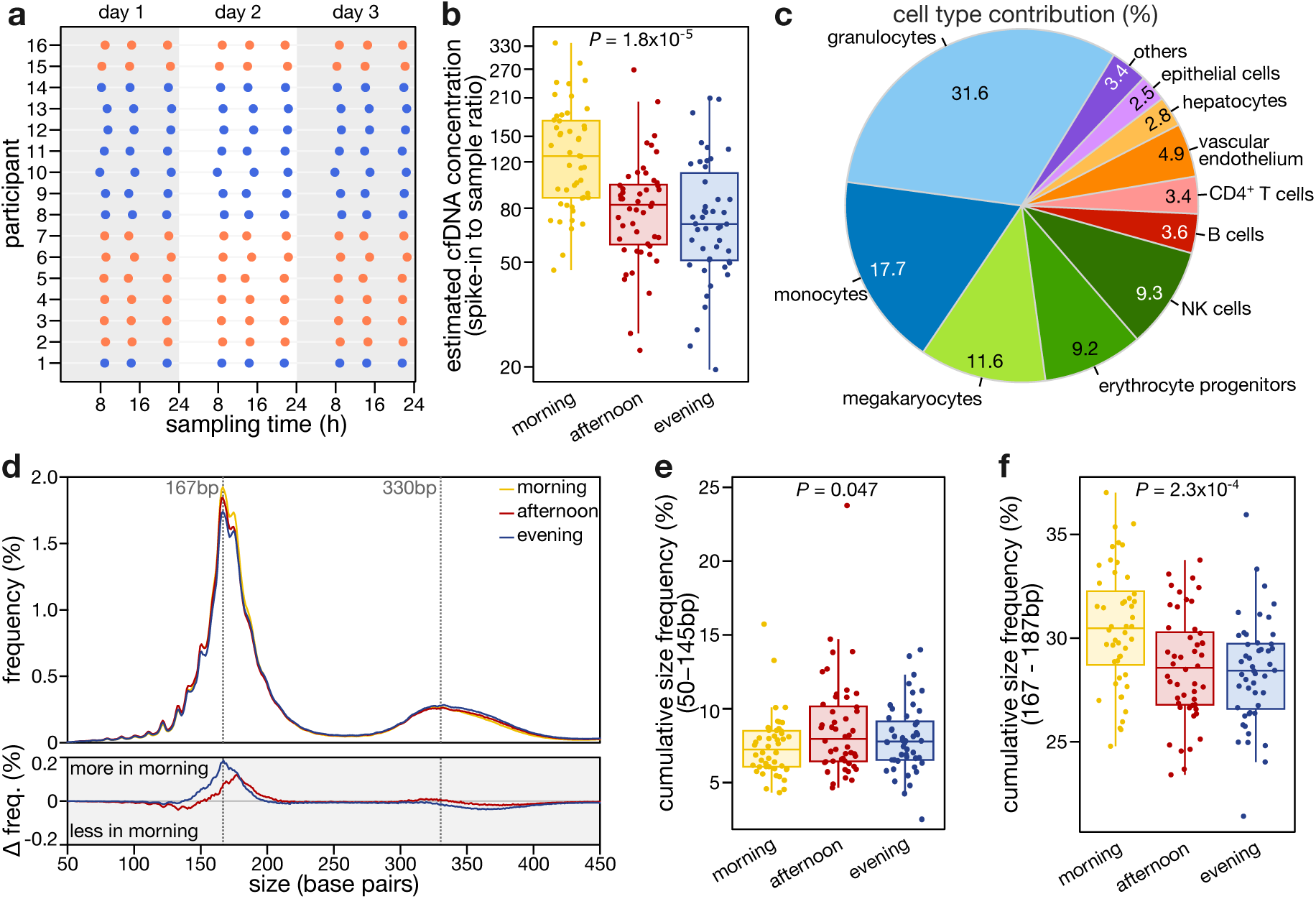
Diurnal variation in cfDNA characteristics. **a**, Overview of diurnal participant sampling across three consecutive days. Sampling time is shown as the interval from midnight in hours for 8 men (blue) and 8 women (orange). **b**, Estimated cfDNA concentration levels at different timepoints: morning, afternoon and evening. **c**, Plasma cell type contributions to cfDNA in the diurnal cohort represented as a pie chart. *Other* cell types include adipocytes, cardiomyocytes, neurons, pancreatic cells, and skeletal muscle cells. **d**, Size distributions of all cfDNA fragments from samples at different timepoints (top) and in morning versus afternoon or evening samples (bottom). **e-f**, Cumulative frequency of subnucleosomal (**e**; 50-145 bp) and mononucleosomal (**f**; 167-187 bp)-sized fragments across timepoints. h: hours. *P* values by repeated measures ANOVA (b, e, f).

We first evaluated if cfDNA concentration estimates vary among individuals and sampling times (Methods; Supplementary Figure 1a). Interestingly, samples collected in the morning were on average 63% more concentrated than in the afternoon and evening (Figure 1b; Supplementary Figure 1b-c; repeated measures ANOVA *P* = 1.8×10^-5^). To investigate if cell types show altered contributions, we quantified the relative fractions of specific cell types in cfDNA through methylation-based deconvolution analysis using EpiDISH^26^ (Methods; Supplementary Figure 1d). In line with earlier reports,^11,12,25^ hematopoietic cell types showed the highest contributions (Figure 1c), with cfDNA of myeloid cell lineage predominating, including granulocytes (31.6%), monocytes (17.7%), megakaryocytes (11.6%) and erythrocyte progenitors (9.2%), followed by cfDNA of lymphoid cell lineage such as B, CD4+ T and natural killer (NK) cells (16.3%). Other large organs with substantial cellular turnover^27^ also showed notable contributions, including vascular endothelium (4.9%), hepatocytes (2.8%) and other epithelial cells (2.5%). Cell type contributions did not differ significantly between sampling times, although proportions of NK cells and hepatocytes tended to be higher and lower in the morning, respectively (Supplementary Figure 1e-f).

Fragment size distributions at different sampling times exhibited typical cfDNA fragmentation profiles, characterized by a dominant peak at ∼167 bp, corresponding to mononucleosomal DNA, and a secondary peak at ∼334 bp, indicative of dinucleosomal DNA. While fragmentation patterns were consistent across sampling times, morning samples showed differences, being more abundant in mononucleosomal while less abundant in subnucleosomal fragments than afternoon or evening samples (Figure 1d). Indeed, a repeated measures ANOVA test showed a significant effect of sampling time on the frequencies of subnucleosomal and mononucleosomal fragments (Figure 1e-f; Supplementary Figure 1g-h; *P =* 0.047 and 2.3×10^-4^). Together, these observations suggest diurnal variation in cfDNA, mostly in concentration but also in fragmentation.

### Demographic variation in cfDNA concentration

We next investigated if demographic factors in addition to sampling time contribute to normal cfDNA variation. We assembled a cohort of 144 adults evenly spread across age ranges, with a similar number of males and females, and all having a normal self-reported health status (Figure 2a; Table 1). We only excluded recent vigorous exercise^28^, and not variables such as high blood pressure or intake of medications such as oral contraceptives, which are common in our medicalized society. Blood plasma from participants was collected by venipuncture at baseline (year 1) and again one year later (year 2) (Supplementary Table 1), resulting in 288 samples. We first assessed variation in cfDNA concentrations cross-sectionally. At both sampling times, cfDNA concentrations exhibited a similar, right-skewed distribution, with an extended tail towards elevated cfDNA levels (Figure 2b). Two individuals with consistent high total cfDNA levels took allopurinol, a xanthine oxidase inhibitor in the purine catabolism that may affect cfDNA turnover. Overall, concentrations of samples from the same individual obtained in both years were significantly correlated (Figure 2c; R = 0.52, *P* = 1.8×10^-11^), and intraindividual variability was notably lower than interindividual variability. The observed higher interindividual variability in part related to differences in sex and BMI, with higher cfDNA concentrations in males (*P* = 3.6×10^-5^, Wilcoxon rank-sum test) and in individuals with higher BMI in line with earlier reports^29–34^ (Figure 2d-e; Supplementary Figure 2a-b; R = 0.27, *P* < 0.001). In contrast to some earlier reports^30,35,36^, we did not observe elevated cfDNA concentrations with increasing age (Supplementary Figure 2c). Finally, leveraging the repeated measurements, we applied a mixed effects model, assessing random and fixed effects of various factors on cfDNA concentration. This confirmed that interindividual differences account for approximately 40% of the total variance in estimated cfDNA concentration, while sampling time and year random effects contribute negligibly (0%) to the total variance. The remaining 60% variance represents residual variability. As this cohort was entirely sampled during daytime, sampling time was indeed unlikely to affect concentrations. From the fixed effect variables, sex and BMI showed a statistically significant association with cfDNA concentration (*P* = 5.7×10^-3^ and 9.8×10^-4^, respectively), while participant age was again not significant (*P* = 1.0; Supplementary Table 2).

**Figure 2:**
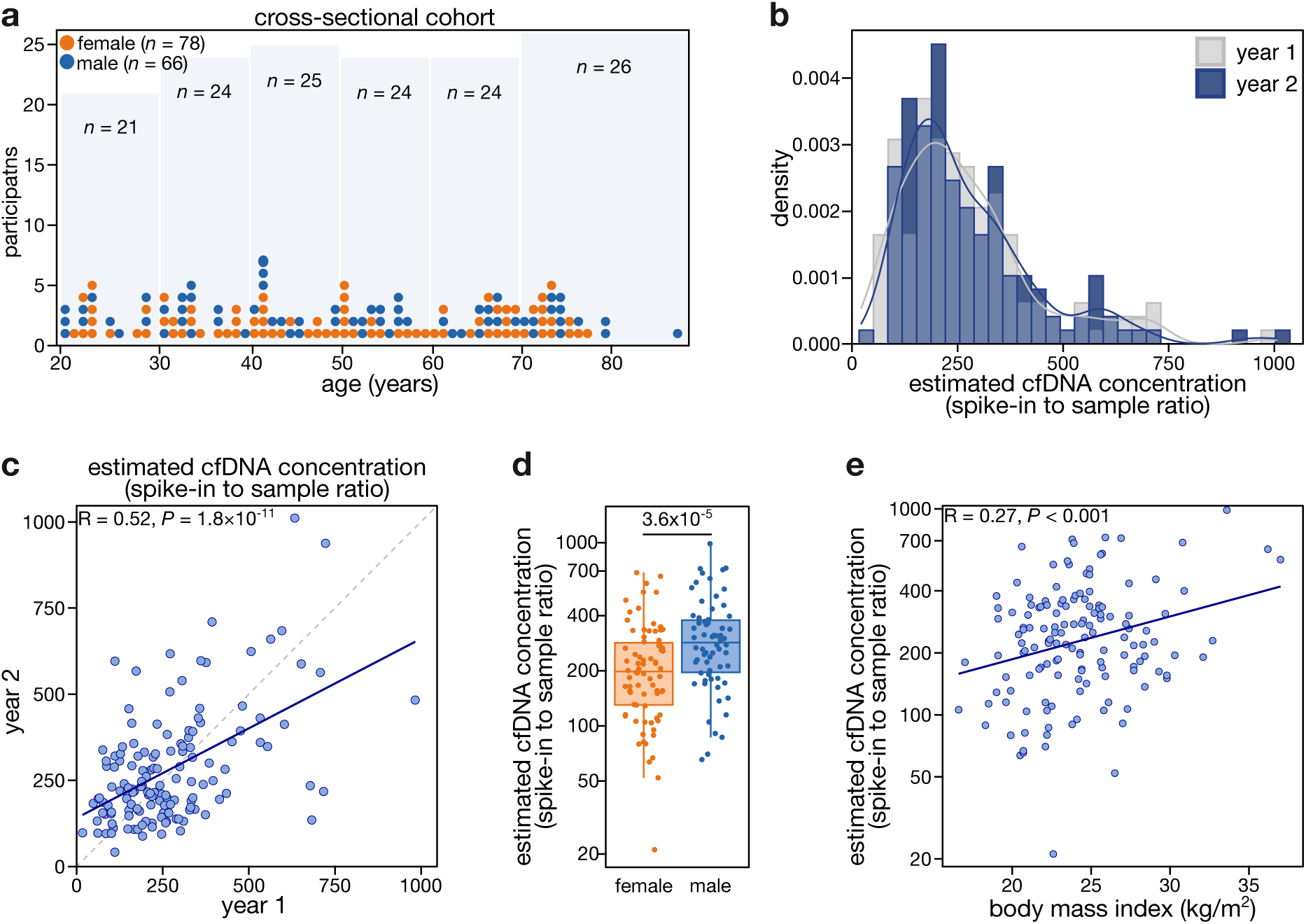
Demographic factors influencing cfDNA concentration. **a**, Overview of sex and age distribution in the cross-sectional cohort (*n* = 144; 78 female, 66 male). **b**, Distribution of estimated cfDNA concentrations at both sampling timepoints. **c**, Pearson correlation of estimated cfDNA concentrations within individuals across the two timepoints. Concentration is expressed as the spike-in to sample ratio. **d**, Comparison of estimated cfDNA concentrations (log10 scaled) between female and male participants in year 1. **e**, Pearson correlation between body mass index and estimated cfDNA concentration (log10 scaled) in year 1. *P* values by Pearson correlation (c, e) or Wilcoxon rank-sum test (d).

### Variation in cfDNA composition

We next assessed relative cell type contributions to plasma cfDNA by deconvolution analyses, first focusing on the samples from year 1. Consistent with observations from the diurnal cohort, hematopoietic cells dominated the cfDNA pool, and also contributions from other cell types showed similar patterns (Figure 3a). We measured complete blood counts for a limited set of samples (*n* = 16). In line with previous reports^37,38^, proportions of blood cell types measured in cfDNA and in whole blood did not correlate (Supplementary Figure 3a). Notably, monocytes showed higher proportions in cfDNA than whole blood (Supplementary Figure 3b; *P* = 1.6×10^-6^, Wilcoxon signed-rank test).

**Figure 3:**
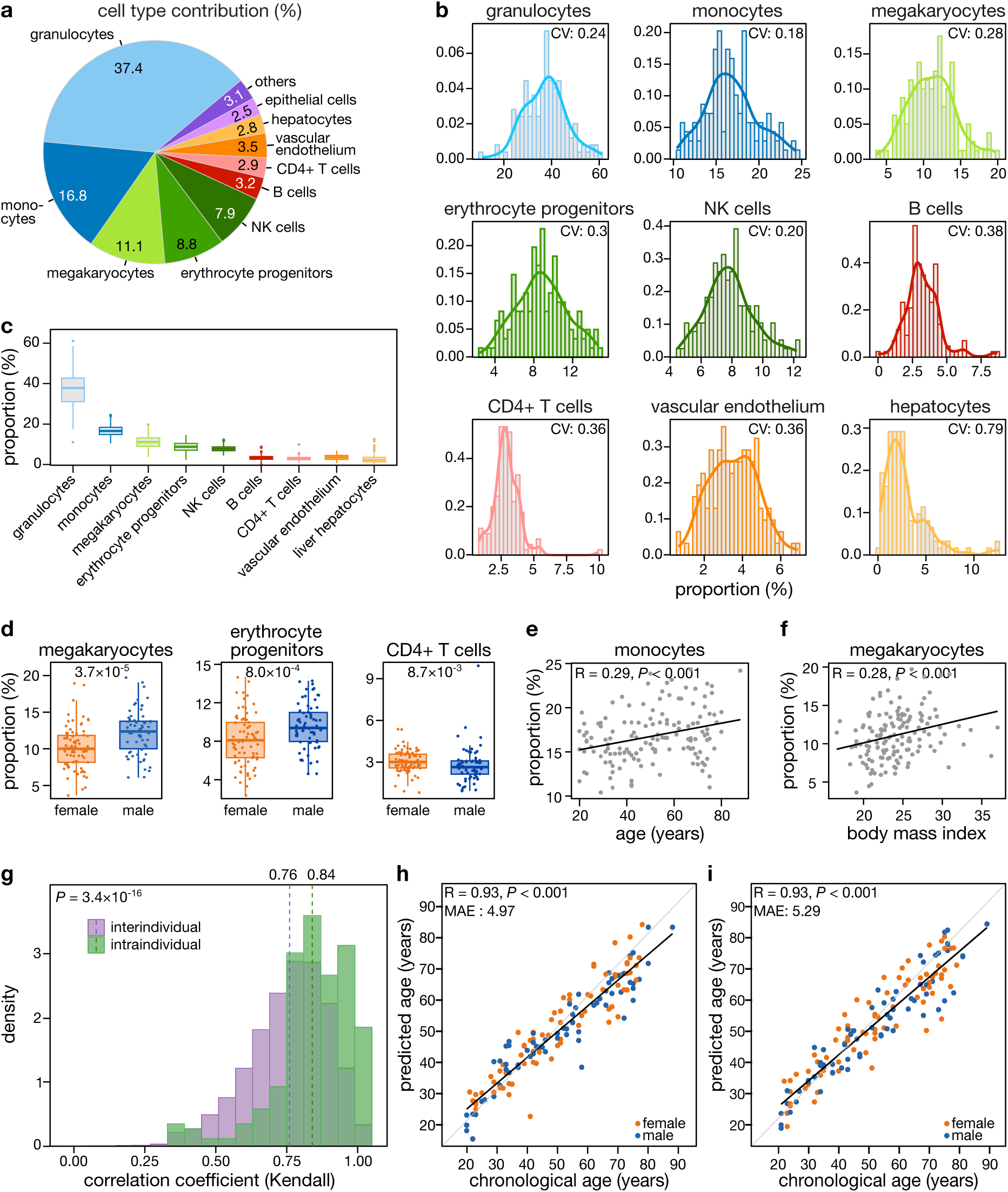
Composition of plasma cfDNA and cfDNAme-based age prediction. **a**, Plasma cell type contributions to cfDNA in year 1 samples, represented as a pie chart. Other cell types include adipocytes, cardiomyocytes, neurons, pancreatic cells, and skeletal muscle cells. **b**, Distribution of proportions (density) for major cell types contributing to plasma cfDNA in year 1. All, except hepatocytes, approximate a normal distribution. **c**, Interindividual variation in major cell types contributing to plasma cfDNA in year 1. **d**, Comparison of cell type contributions to plasma cfDNA between female and male participants in year 1. **e**, Pearson correlation between monocyte proportions and age in year 1 samples. **f**, Pearson correlation between megakaryocyte proportions and body mass index (kg/m^2^) year 1 samples. **g**, Kendall correlation coefficients showing inter- and intraindividual variability in cell type composition across both sampling timepoints for major contributing cell types. **h**, Performance of an elastic-net regression model for cfDNA methylation-based age prediction, trained on the variable sex and 146 CpGs using year 1 samples. **i**, Performance of the age prediction model in the validation set (year 2 samples). CV: coefficient of variation. MAE: mean absolute error. *P* values by Pearson correlation (e, f, h, i) or Wilcoxon rank-sum test (d, g).

The inferred proportions of most major cell types approximated a normal distribution (Figure 3b). However, the degree of interindividual variability varied by cell type (Figure 3c; Supplementary Table 3). The highest coefficient of variation (coefficient of variation CV = 0.79) was observed for hepatocytes, which also showed a right-skewed distribution. Interestingly, several demographic variables correlated with the contribution of specific cell types. Megakaryocytes and erythrocyte progenitor cells were elevated in males versus females, while CD4+ T cells were lower (Figure 3d, *P* = 3.7×10^-5^, 8.0×10^-4^ and 8.7×10^-3^, Wilcoxon rank-sum test). We also observed higher monocyte contributions with increasing age, and higher megakaryocyte contributions with increasing BMI (Figure 3e-f). Despite notable interindividual variability, correlations were observed between cell type fractions. A negative correlation between granulocytes and most other cell types (Supplementary Figure 3c), while a positive correlation between megakaryocytes and erythrocyte progenitors was detected (Pearson R = 0.89, *P* = 2.0×10^-49^).

We next verified if our observations from year 1 samples were replicated in year 2. The immune cell type fractions exhibited similar correlations, proportions, distributional patterns and variability across the two timepoints (Supplementary Figure 3c-f; Supplementary Table 3). Sex-specific differences in the proportions of megakaryocytes, erythrocyte progenitors and CD4+ T cells were consistently observed, as were the associations of age with monocytes and BMI with megakaryocytes (Supplementary Figure 3g-i). To further confirm these associations, we constructed linear mixed-effect models which supported the significance of these relationships (Supplementary Table 4). Analysis of the random effects revealed that the contributions of megakaryocytes were stable within the same individual across both years (intra-class correlation coefficient, ICC = 0.767), whereas those of B and CD4+ T cells were more variable (ICC = 0.1514 and 0.2662, respectively). Finally, we compared intraindividual and interindividual variation in cell type compositions by quantifying Kendall’s correlation coefficients for major contributing cell types between individuals and over time. Here, the intraindividual correlation was significantly higher than the interindividual correlation (Figure 3g; *P* = 3.4×10^-16^, Wilcoxon rank-sum test). The contribution of different cell types to cfDNA is thus relatively stable within an individual over this time window.

### cfDNA methylation-based age prediction

Various studies have explored cfDNAme changes in aging^39,40^, but data on their predictive ability is scarce. While we observed a statistically significant correlation between monocytes and age (R = 0.29), this modest association alone was insufficient for a reliable predictive model. To enable cfDNAme-based age estimation, we developed a cfDNAme-based age prediction model. From the year 1 dataset, we first identified age-correlated CpGs from the capture panel by selecting sites with an absolute Spearman correlation coefficient greater than 0.1, yielding 9,275 candidate features. We then trained a penalized elastic net regression model, incorporating sex as a non-penalized covariate. Adopting a leave-one-out (LOO) cross-validation strategy yielded 146 CpGs in the final model (Supplementary Table 5). Two of these CpGs showed T-cell-specific methylation, and many (53/146) were in the vicinity of CpGs previously used in age prediction models^41,42^. Applying this model on the training set (year 1) produced accurate age estimates, with a mean absolute error (MAE) of 4 years and 11 months (Figure 3h; Pearson R = 0.93, *P* < 0.001). Importantly, validation of this model on the year 2 dataset yielded a comparable MAE of 5 years, 3 months (Figure 3i; Pearson R = 0.93, *P* < 0.001), indicating that our model can predict chronological age from cfDNAme data.

### cfDNA fragmentation patterns

We next assessed cfDNA fragmentation patterns. Here, fragment size profiles (50-450 bp) showed modestly higher intraindividual *versus* interindividual correlations (mean Kendall’s correlation = 0.79 vs 0.78; *P* = 0.13, Wilcox rank-sum test). Abundant cfDNA sizes reflecting mono- and dinucleosomal fragments were present at more variable rates (standard deviation SD = 0.15) than less abundant sizes reflecting subnucleosomal fragments (SD = 0.05). Sex, age, and BMI were however not significantly associated with fragment size patterns in this cross-sectional cohort (Figure 4a; Supplementary Figure 4a). Notably, some size ranges within our cohort were strongly correlated with one another. For example, mononucleosomal fragments (167-187 bp) were positively correlated with subnucleosomal fragments (<145 bp) but negatively correlated with dinucleosomal fragments (330-420 bp) (Figure 4b-d). These correlations were replicated in year 2 samples (Supplementary Figure 4b-d), highlighting a structured fragmentation landscape. These sets of correlated fragment lengths determined fragment size bins that were used throughout this study. The opposing abundance patterns suggest a discrete biogenesis, potentially reflecting differences in nuclease activity, cell death pathways and nucleosome positioning.

**Figure 4:**
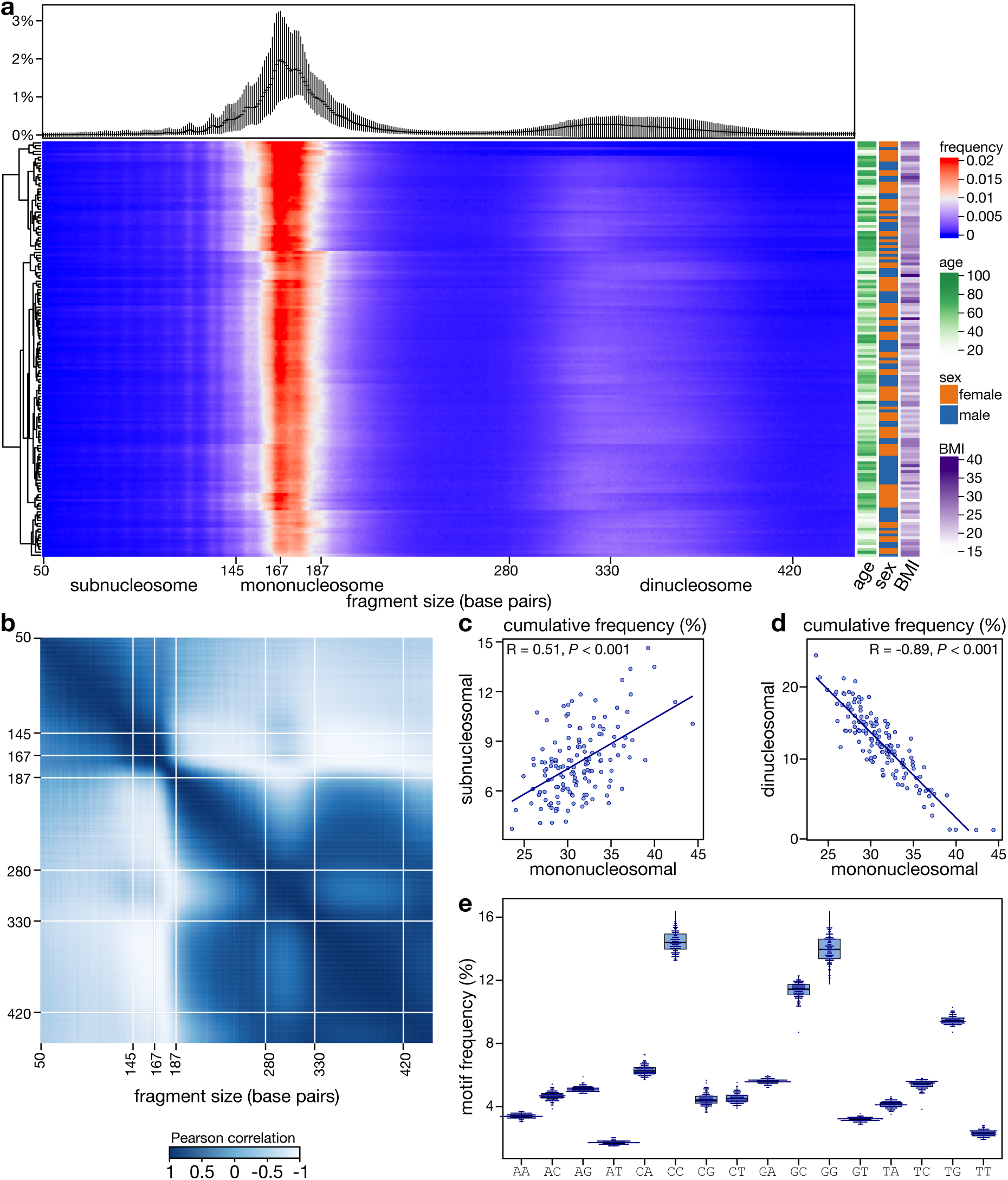
cfDNA fragmentation patterns in the cross-sectional cohort. **a**, cfDNA fragment size distributions in year 1 samples, shown as average profiles with error bars representing upper and lower quantiles (top), and a correlation matrix (Pearson) showing associations between fragment sizes and demographic variables (sex, age, and BMI; bottom right). Fragment sizes are defined as subnucleosomal (<145 bp), mononucleosomal (167-187 bp) and dinucleosomal (330-420 bp). **b**, Correlations of fragment sizes in year 1 samples. **c**,**d**, correlation between mononucleosomal and subnucleosomal fragments (c) or dinucleosomal fragments (d). **e**, Relative contributions of dinucleotide end motifs to plasma cfDNA in year 1 samples. BMI: body mass index (kg/m²); age in years. *P* values by Pearson correlation (c, d).

In addition to fragment size, we also assessed dinucleotide end motifs (2-mer sequences). Note that while the nature of targeted sequencing may affect sequence end context, we reason that this effect is systematic across all samples. The CC motif was most abundant, with an average frequency of 14.4% in our cohort (Figure 4e; Supplementary Figure 4e), consistent with prior studies^22,43,44^. Intraindividual correlations of end motifs were higher than interindividual correlations (mean Kendall’s correlation = 0.94 versus 0.93; *P* = 4.6×10^-3^, Wilcox rank-sum test), and specific end motifs were associated with sex, age, and BMI (Supplementary Figure 5a-c). Motif-motif correlation revealed A/T-rich motifs were largely positively correlated among themselves. C-end motifs tended to negatively correlate with G- end motifs, with CG as an exception (Supplementary Figure 5d-e).

### Distinct processes underlying cfDNA fragmentation

The data generated enables us to not only assess the impact of demographic variables on the make-up of cfDNA, it also provides a window into the processes underlying cfDNA biogenesis. We first assessed if some cfDNA fragment sizes have specific end motifs, possibly reflecting the activity of different nucleases. We therefore calculated the frequencies of fragment sizes and 2-mer end motifs in each of the 144 samples and assessed the correlation between both. This revealed both positive and negative associations, indicating that some end motifs are enriched in fragments of certain size ranges, with consistent correlations emerging for either subnucleosomal, mononucleosomal or dinucleosomal fragments. These correlations were evident in samples from both time points (Figure 5a; Supplementary Figure 6a). For example, C-end motifs tended to be more abundant in shorter fragments, while G-end motifs were more common in longer fragments. The CC motif, in particular, was enriched within mononucleosomal fragments (167-187 bp). Complementing the size-size and motif-motif correlations — which revealed coordinated abundance shifts around mononucleosomal lengths and a negative association between C- and G-end motifs, with their preferential enrichment in short and long fragments respectively — this analysis provides additional insights into cfDNA fragmentation patterns. Together, these data suggest that fragment size and their associated end motifs likely reflect the activity of distinct sets of nucleases acting under specific chromatin or cellular environments, contributing differentially to the generation of shorter or longer fragments.

**Figure 5:**
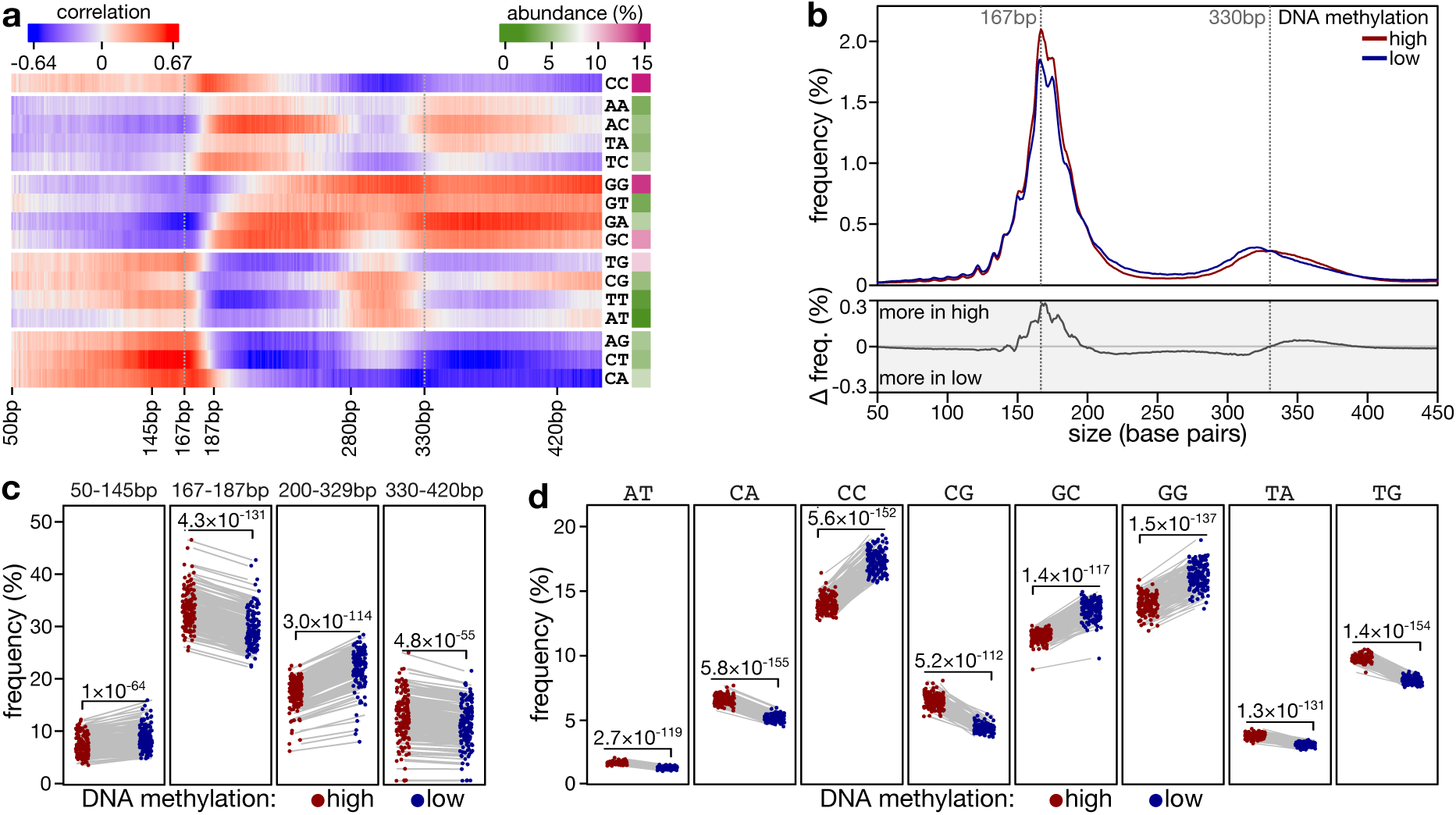
Factors associated with cfDNA fragmentome characteristics. **a**, Correlation matrix (Spearman) showing associations between cfDNA fragment sizes (in bp resolution) and 2-mer end motifs in year 1 samples, for subnucleosomal (<145 bp), mononucleosomal (167-187 bp), and dinucleosomal fragments (330-420 bp). **b**, Fragment size distributions (top) and frequency difference (bottom) of cfDNA fragments overlapping CpGs that are either highly methylated (red; *n* = 2,953) or lowly methylated (blue; *n* = 2,752) across major contributing cell types in year 1 (Methods). **c**, Cumulative frequency of highly and lowly methylated fragments across different fragment size ranges in year 1 samples. **d**, Frequency of highly and lowly methylated fragments across different 2-mer end motifs in year 1 samples. bp: base pairs. *P* values by paired t-test (c, d).

In addition to nuclease activity, chromatin compaction and other epigenetic features may also influence fragmentation characteristics. The interplay between these factors remains underexplored and a deeper understanding is key to maximize exploiting liquid biopsies. We therefore first assessed the correlation between DNAme and cfDNA fragmentation: low DNAme levels characterize accessible chromatin, while DNA in compacted chromatin is typically highly methylated. To study this, we selected CpGs that were either highly methylated (*n* = 2,953) or lowly methylated (*n* = 2,752) across the major cfDNA-contributing cell types (Methods). Unsurprisingly, these CpGs retained their respective methylation states in cfDNA, allowing analysis of fragment characteristics in different methylation contexts (Supplementary Figure 6b). We then analysed all fragments that overlap these CpGs. Fragments overlapping highly methylated CpGs were more enriched in mono- and dinucleosomal sizes (*P* = 4.2×10^-131^ and 4.7×10^-55^, paired t-test), while fragments derived from lowly methylated regions were more abundant in subnucleosomal fragments (*P* = 1.0×10^-64^, paired t-test), in line with these fragments deriving from less defined, nucleosome-depleted chromatin (Figure 5b-c; Supplementary Figure 6c-d)^2,45^.

End motif analysis further highlighted differential cleavage patterns between both sets of fragments. Specifically, CC, GC and GG motifs were particular enriched in low methylation fragments (all *P* < 1.0×10^-116^, paired t-test), and CA, CG and others in high methylation fragments (Figure 5d; Supplementary Figure 6e all *P* < 1.0×10^-111^, paired t-test). The CG motif reflects the canonical DNA methylation site, and its enrichment in methylated fragments is in line with earlier studies demonstrating that DNASE1L3, the leading endonuclease in plasma, has a strong preference for cleavage at methylated cytosines^23^. Additionally, in our data, the frequency of CG motif cleavage at highly methylated sites was negatively correlated with estimated cfDNA concentration (Supplementary Figure 6f).

The CC motif, the most abundant 2-mer end motif and corresponding to the preferred DNASE1L3 cutting site^43^, was more frequent in lowly methylated regions. The reduction of CC and elevation of CG and CA cleavage in highly methylated contexts suggests that epigenetic states influence the activity or preference of some nucleases, and of DNASE1L3 in particular, in plasma cfDNA. Furthermore, the reduced abundance of mononucleosomal-sized fragments in low methylation regions likely reflects a higher probability of cleavage in these more accessible regions. Collectively, these data indicate that the epigenome affects cfDNA cleavage and that CpG methylation modulates nuclease activity.

### Cell-type-specific effects on cfDNA fragmentation

An outstanding question is whether cfDNA from every cell type is processed similarly, or whether the cellular origin of cfDNA is also reflected in its general fragmentation pattern. To address this, we first investigated associations between the cell type compositions of cfDNA and its fragmentation pattern. With cfDNA fragment size and end motif intricately correlated, we jointly analysed both modalities in relation to contributing cell types. This revealed a strong association with granulocytes: samples with higher granulocyte contributions showed a reduced fraction of subnucleosomal fragments and a pronounced enrichment of mononucleosomal fragments, particularly for fragments of ∼167 bp and longer (Figure 6a; Supplementary Figure 7a). This pattern was further illustrated by directly comparing the ten samples with the highest versus the lowest granulocyte fractions. Here, samples with a higher granulocyte contribution had fewer short fragments (<166 bp; Figure 6b; Supplementary Figure 7b). Also other cell types correlated with fragment size and end (Figure 6a; Supplementary Figure 7a). While the broad trends in fragmentation appeared consistent across different end motifs, we did observe variation between cell types. These observations support the idea that cfDNA fragmentation is not uniform across all cell types and that intrinsic cellular properties, such as chromatin structure, epigenetic state, and nuclease activity, may influence the fragmentation in a cell-type-specific manner.

**Figure 6:**
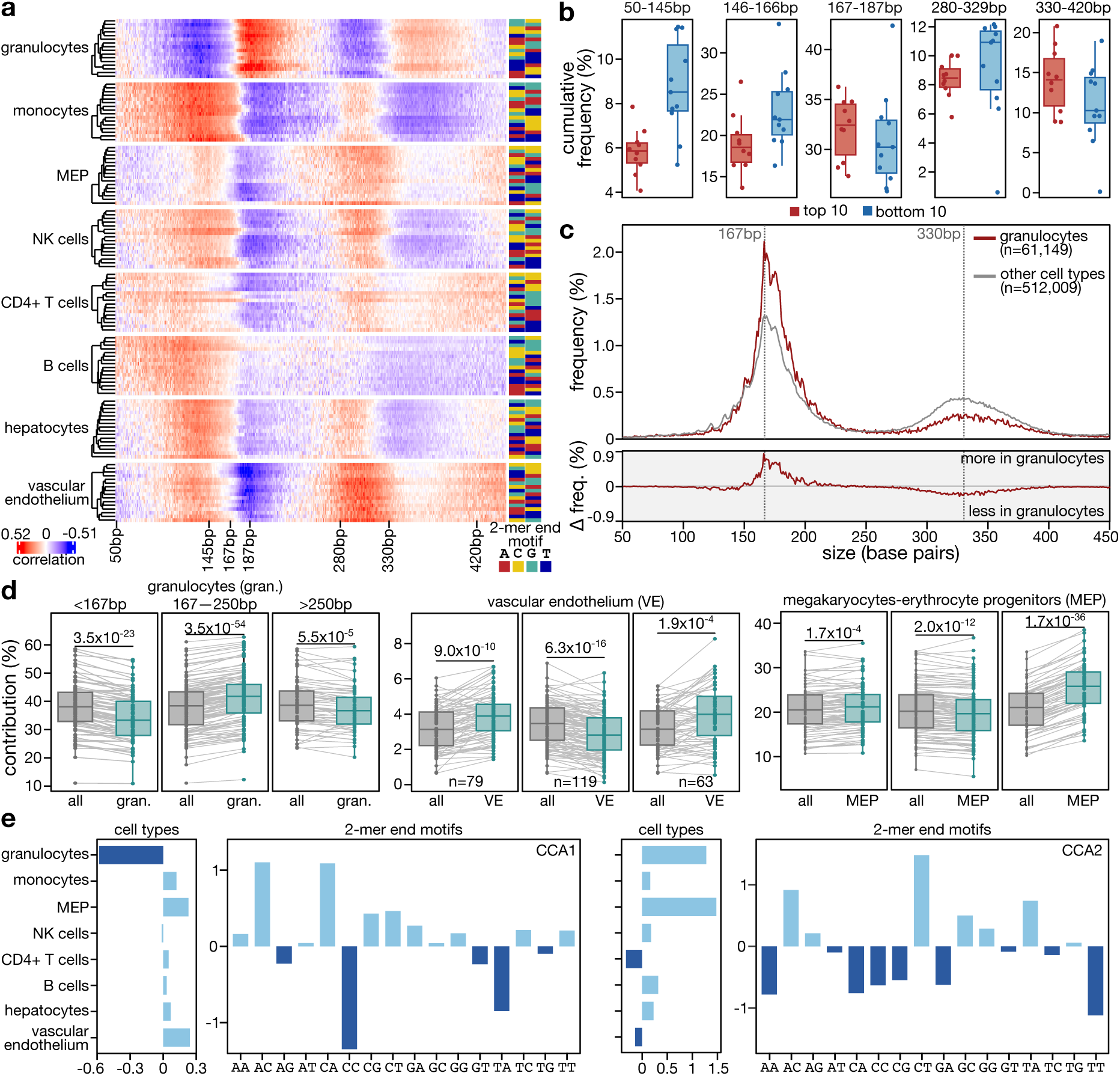
Cell-type-specific effects on cfDNA fragmentation. **a**, Correlation matrix (Spearman) showing cell-type specific associations between cfDNA fragment sizes and 2-mer end motifs for major contributing cell types in year 1 samples. **b**, Cumulative frequency of fragments across different fragment size ranges in the top ten samples with the highest versus lowest granulocyte proportions in year 1. **c**, Fragment size distributions (top) and frequency difference (bottom) between hypomethylated granulocyte-specific fragments and all other hypomethylated cell type-specific fragments in year 1 samples. Fragment selection based on cell type-specific markers with >60% methylation difference. **d**, Deconvolution analysis of cfDNA from year 1 samples after *in silico* size selection into three bins (<167 bp, 167–250 bp, >250 bp); samples with <10× coverage per bin were excluded, retaining 79, 119, and 63 samples per bin respectively. **e**, Canonical correlation analysis (CCA) showing the association between cell type composition and cfDNA 2-mer end motifs in the first two dimensions. bp: base pairs; MEP: megakaryocytes-erythrocyte progenitors; VE: vascular endothelium. *P* values by paired t-test (d).

To more directly assess the relationship between cell type and fragment size, we leveraged cell-type-specific DNA methylation markers: we selected a total of 528 hypomethylated (233 regions) and 282 hypermethylated (92 regions) markers, where average DNA methylation levels for that cell type differed by >60% from other types (Supplementary Figure 7c; Supplementary Table 6). We next parsed reads overlapping these CpGs and stratified them based on their DNA methylation status (methylated or unmethylated), assigning fragments to the corresponding cell type. Due to missing or low numbers of hypermethylated markers across the major cell types except for hepatocytes, we focused our analysis on fragments derived from hypomethylated markers. This analysis was performed by pooling all samples from year 1, allowing us to obtain enough reads for every cell type. Granulocyte-specific fragments exhibited an enrichment in mononucleosomal sizes, echoing our prior correlation-based analysis linking higher granulocyte contributions with mononucleosomal fragments (Figure 6c; Supplementary Figure 7d). Although hepatocyte-specific fragments displayed a distinct size profile, with a depletion in mononucleosomal fragments and a relative enrichment in both subnucleosomal and dinucleosomal sizes compared to fragments from other cell types over hypomethylated markers, this was not observed over hypermethylated markers (Supplementary Figure 7e). Hepatocyte-specific fragments derived from hypomethylated markers tended to be shorter comparing to other cell types, which was also evident in the correlation analysis. The discrepancies may stem from differences in sensitivity and resolution of the analyses, in which marker-based classification relies on more specific, but sparser, CpG-defined assignments. Together, these data suggest that cfDNA fragments from specific cell types show characteristic size profiles, and that certain cell types contribute to specific cfDNA size ranges in a disproportionate manner. This was further demonstrated through *in silico* size selection, binning cfDNA into three ranges (<167 bp; 167-250 bp and >250 bp), and performing deconvolution analysis on these bins. To ensure robustness, we excluded samples with a coverage <10× per bin, retaining 79, 119, and 63 out 144 samples from year 1 for analysis (Methods). As expected, the granulocytes contribution was reduced in short and in long fragment bins (<167 bp and >250 bp) but increased in the mononucleosomal fragment bin (167-250 bp), while contributions from megakaryocytes-erythrocyte progenitors (MEP) and vascular endothelium (VE) cells were increased in short (<167 bp) and long fragments (>250 bp) (Figure 6d; Supplementary Figure 7f).

Finally, to further investigate the association between cell type fractions and end motif, we performed regularized canonical correlation analysis (CCA) using all 288 samples. The first 2 canonical correlations (0.75 and 0.65 respectively; Supplementary Figure 7g) suggested that cfDNA fragmentation was strongly linked to cell type composition (Supplementary Figure 7h; permutation test *P* = 0). We examined the canonical loadings to identify cell types and end motif that contributed most to the first two canonical components. In the first component, granulocytes showed the strongest negative loading on CCA1, followed by positive contributions from vascular endothelium and megakaryocytes-erythrocyte progenitors. For this component, the CC motif exhibited the strongest negative loadings while the AC and CA motifs showed strong positive loadings (Figure 6e). Together, these data suggest that the granulocyte contribution tends to be associated with the CC motif, while vascular endothelium and megakaryocytes-erythrocyte progenitors are likely linked to the CA and AC motifs.

## DISCUSSION

While cfDNA-based liquid biopsies increasingly gain traction for diagnostics, prognostics and therapy monitoring, variables affecting the baseline characteristics of plasma-derived cfDNA in the healthy population, including composition, concentration and fragmentation, remain incompletely understood. Here, we studied cfDNA from 432 plasma samples of healthy individuals using targeted enzymatic methyl-sequencing. We observe that different demography- and sampling-related variables shape cfDNA, and that cfDNA originates from diverse cell types in a dynamic and individual-specific manner. Fragment size and end motifs are only moderately influenced by demographic variables, but appear intricately linked to cell-type-specific effects governed by nucleosome positioning and nuclease activity. Our findings not only inform the design of cfDNA-based clinical studies but also yield insights into the role of specific cell types in cfDNA biogenesis, processing and presentation.

Deconvolution analysis showed that cfDNA was predominantly of hematopoietic origin, as previously established^11,12,25^. Individuals differed in total cfDNA levels and cell-type contributions, but the intraindividual variation was low across samples collected over hours, days or a year. This intraindividual stability likely reflects intrinsic hematopoietic setpoints^38,46,47^. Among the hematopoietic lineages, the granulocyte contribution was particularly variable between individuals. This is reminiscent of the high granulocyte count variability in the healthy population^48^, although blood cell counts did not correlate with their cfDNA contribution, suggesting other factors such as cell turnover, clearance or death mechanisms may be involved. We further assessed the influence of demographic factors, including sex, age, and BMI, on cfDNA characteristics. These factors significantly influenced cfDNA levels and composition: men exhibited higher cfDNA concentrations and increased contributions from megakaryocytes and erythroid progenitors, while women showed more T-cell-derived cfDNA, potentially reflecting sex-specific immune regulation, including estrogen-driven increase in T-cell turnover^49^. Higher BMI correlated with increased cfDNA concentration and megakaryocyte contribution. With advancing age, total cfDNA remained stable, but the monocyte-derived fraction increased. This mirrors previous findings in older populations and may reflect myeloid skewing in hematopoiesis^38,50^. Beyond age-related changes in cell type composition, we also constructed a cfDNAme-based epigenetic clock, achieving a mean absolute error of 5.3 years—comparable to first-generation clocks^41,42^. Another notable observation was that sampling time affected cfDNA properties. Unlike earlier studies^51–55^, we sampled our diurnal cohort throughout the day, on 3 consecutive days, and in the participants’ daily environment. cfDNA was consistently more concentrated in early morning samples. This observation aligns with some but not all other studies^30,51,56,57^, with discrepancies potentially originating from differences in study design, sample size, or timing of blood draws. Together, our findings demonstrate that while cfDNA characteristics are relatively stable within healthy individuals over time, they are influenced by demographic and sampling-related variables. This emphasizes the need to account for demographic variables when analyzing cfDNA, and to ensure matching of sampling and demographic variables in case-control study designs to minimize confounding. On the other hand, the observed differences in cfDNA concentration between early morning and afternoon or evening, can also be used to strengthen studies that require high cfDNA concentrations.

It is generally assumed that cfDNA fragmentation is relatively homogeneous in healthy individuals. Consistent with this, our analysis of fragment size profiles revealed recurrent peaks of mono- and dinucleosomal sizes, reflecting the protection histones confer to chromatinized DNA during apoptotic cleavage. These size profiles were largely invariant across demographic factors, indicating that the core chromatin organization and apoptotic cleavage machinery are tightly regulated in healthy states. In parallel, we observed distinct associations between fragment sizes and the cfDNA dinucleotide end motifs that reflect nuclease cleavage preferences. Some end motifs showed a moderate association with demographic factors, implying that physiological factors can subtly modulate nuclease activity. Notably, end motifs preferentially associated with fragments of specific lengths.

Specifically, fragments bearing C-end were predominately positively associated with shorter fragments, while those with G-end motifs tended to occur in longer fragments. End motifs are not independent of size and may reflect shared upstream processes of cfDNA fragmentation. G-end motifs may thus derive from other endonuclease activities than C-rich motifs, which are linked to DNASE1L3 activity^43^. Of note, remarkable size-motif-related shifts appeared between 167 bp and 187 bp, i.e. the size of a chromatosome complexed with a linker histone. This transition pattern may reflect nucleosome compaction states or variability in linker histone H1 binding. Granulocyte-derived fragments contributed disproportionately to mononucleosomal peaks and specific end motifs. It is tempting to speculate that such changes can be ascribed to the active chromatin release mechanism of neutrophils, such as NETosis^28,58^.

It was postulated that the consistency of size profiles in healthy individuals is attributable to uniform cfDNA degradation dynamics, chromatin structures and the homogeneity of cell-of-origin especially with respect to immune cell types ^59^. While in agreement with other studies that immune cells are the dominant contributors in plasma cfDNA, we observed substantial variation in the proportions of contributing cell types across healthy individuals, particularly granulocytes. This implies that stable cfDNA size profiles likely stem from chromatin organization rather than homogeneous cell-of-origin profiles. Furthermore, we observed cell type-specific fragmentation footprints. Granulocyte-derived cfDNA consistently exhibited an enrichment of mononucleosomal fragments across analytical approaches and systematically diverged from other hematopoietic and tissue-derived lineages. Previous studies already demonstrated that cfDNA fragments of non-hematopoietic origin, such as those derived from tumors or from placentas, are typically shorter. Implementing size selection during cfDNA library preparation may inadvertently affect the cell composition of cfDNA^60–62^. Vascular endothelium was associated with enrichment in intermediate sized fragments, both in the correlation analysis and following *in silico* size selection. Hepatocyte-specific fragments appeared to be shorter. Although both technical constraints and differences in the nature of the signal precluded a 1-on-1 comparison with results obtained from our marker-based fragment parsing, these findings collectively reinforce size profiling as an analytical axis of cfDNA, indicating that size selection can be strategically applied to enrich or deplete cfDNA derived from specific tissues. We in addition show that end motifs are influenced by cell types, with the abundant CC-motif associated with granulocytes. Together, these insights suggest that leveraging both size and motif signatures to selectively enrich cfDNA from tissue of interest, stands to enhance diagnostic sensitivities.

We observed a notable negative correlation between granulocytes and other hematopoietic lineages, particularly MEP, across healthy individuals. This reciprocal pattern was mirrored in fragmentation characteristics, in which hematopoietic cells other than granulocytes negatively correlated with mononucleosomal fragments and show distinct motif patterns. This may be partly explained by granulocytes being the main contributors, producing a relative decline of other cell type contribution when they are particularly abundant. Another explanation lies in differential exposure or susceptibility to nuclease activity during cfDNA generation. Supporting this, recent evidence shows that DNA from megakaryocytes remains resistant to DNase treatment in platelet preparations, while leukocyte- and hepatocyte-derived DNA is readily degraded under the same conditions^63^. This suggests that megakaryocyte DNA may be structurally shielded from some nucleases. Additionally, these contrasting profiles may originate from lineage-specific chromatin organization, nuclease expression or cell death pathways.

Shortening of fragment sizes and changes in the frequency of end motifs were observed in hypomethylated cfDNA fragments in line with earlier studies^24,64^. These changes suggest that the DNA methylation status can influence nuclease accessibility or cleavage efficiency, embedding fragmentation within a broader epigenetic regulatory context. In this light, fragmentation characteristics not only mirror cell-type composition but also reflect the underlying chromatin state. Current deconvolution methods are fragmentation-agnostic. Although they remain broadly valid, particularly for cell types with high contribution, the observed interdependencies between cfDNA fragmentation and methylation have important implications for cfDNA methylation-based deconvolution, and fragmentation-aware deconvolution models that tackle cleavage biases stand to further improve resolution and accuracy for tissue-of-origin inference of cfDNA.

In conclusion, our findings support the view that cfDNA is a biologically coordinated system in which chromatin structure, DNA methylation, and nuclease activity jointly define its molecular features. Future work that targets pivotal cell types across diverse physiological and disease contexts is warranted. Moreover, we anticipate that a thorough assessment of existing markers and the discovery of new markers that leverage cfDNA signals in a cell-specific manner will further elucidate cfDNA biogenesis as well improve clinical utility.

## MATERIALS & METHODS

### Recruitment and sampling

The study (no. S66450) was approved by the Medical Ethics Committee of University Hospitals Leuven.

#### a. Cross-sectional cohort

*This cohort compromised 150 participants who were initially sampled at year 1, with a subsequent sampling conducted one year later at year 2*.

Participants were recruited and selected at the Leuven University Hospital and through on-campus initiatives. They were invited to the Leuven University Hospital for peripheral blood sampling, which was conducted using Cell-Free DNA BCT tubes (Streck) following informed consent. All were of European ancestry, except for one Asian participant. An even distribution across 10-year age intervals, ranging from 20 to >70 years, and between male and female participants was targeted. Participants also completed a detailed questionnaire covering demographic information, biometric data, relevant medical history, and lifestyle factors.

Inclusion criteria required participants to report no active or chronic diseases and no ongoing pregnancy. None of the participants reported high action sports within one hour prior to sampling. Disease status was based on self-assessment. Initial sample collection for year 1 (*n* = 150) was conducted between October 2022 and March 2023. At a follow-up invitation, five participants were lost to follow-up, resulting in 145 samples for year 2. Samples for year 2 were collected one year after the first visit. Efforts were made to schedule the second visit as close to exactly one year later and at the same time of day (Supplementary Table 1). Identical blood collection tubes were used at both timepoints to ensure consistency.

#### b. Diurnal cohort

*This cohort comprised 16 participants (equal male and female) who underwent diurnal sampling in the morning, afternoon, and evening over three consecutive days*.

Participants were recruited through on-campus announcement, and following an initial health assessment, peripheral venipuncture was performed at nine timepoints using Cell3™ Preserver tubes (Nonacus). Samples were collected in the participants’ usual environments at defined intervals: morning (7:50–9:30 am), afternoon (1:30–3:20 pm), and evening (9:30–10:35 pm) (Supplementary Table 1). Participants were instructed to maintain their usual daily activities to maximize generalizability except for heavy exercise prior to sampling. Timing of diurnal sampling for each participant varied by no more than 30 minutes across corresponding timepoints on different days (Figure 1a). Sampling was conducted between April and June 2023.

### Sample processing

Samples were kept and transported at room temperature prior to plasma separation. Plasma separation was performed using a double spin protocol according to manufacturer’s instructions. For the cross-sectional cohort, plasma separation was performed within one week after sampling. For the diurnal cohort, separation of a participant’s samples was performed the day following the last sampling day. Following double centrifugation, the recovered plasma (approx. 4-5ml) was aliquoted in 2 ml LoBind tubes (Eppendorf) and stored at -20°C or -80°C. cfDNA extraction was performed automatically from 2 mL of plasma using the Maxwell® RSC ccfDNA Plasma Kit (Promega) on the Hamilton Liquid Handler following standardized protocols.

Complete blood counts (*n* = 16) were measured by the Advia 2120 hemocytometer following manufacturer’s instructions.

### Library preparation

For library generation, 18 µL of extracted cfDNA per sample was used. Fixed quantities of sonicated unmethylated lambda DNA and CpG methylated pUC DNA were spiked into each cfDNA sample prior to EM-seq library construction to evaluate library conversion efficiency and allow concentration estimation. Libraries were generated using Enzymatic Methyl-seq (EM-seq, New England Biolabs) following an in-house optimized protocol with half the recommended reagent volumes, a 30-minute adapter ligation step instead of the standard 15 minutes and 12 polymerase chain reaction cycles. Library concentrations were measured using a Nanodrop spectrophotometer and fragment profiles were assessed using Bioanalyzer HS (High Sensitivity DNA Kit, Agilent) or Fragment Analyzer 5200 (HS NGS Fragment Kit (1-6000bp), Agilent).

### Capture-based target enrichment and sequencing

Processed libraries were pooled in equal mass amounts and subjected to capture-based target enrichment using a custom developed probe set covering 4,991 regions showing cell-specific methylation for tissue and cell types that possibly contribute to cfDNA, including hematopoietic cells and solid tissue types (Roche KAPA HyperExplore Probes).

Target enrichment was performed as previously described^65^, and the resulting libraries were sequenced on Novaseq instruments (Illumina) in 150-bp paired-end mode. Sequencing data were uniformly processed by the nf-core/methylseq pipeline^66^ (2.6.0). Reads were aligned to hg38 using bismark (0.24.0) without hardclipping. Specific parameters were set as below.

clip_r1: 0

clip_r2: 0

three_prime_clip_r1: 0

three_prime_clip_r2: 0

aligner: ‘bismark’

ignore_r1: 10

ignore_r2: 7

ignore_3prime_r1: 15

ignore_3prime_r2: 13

meth_cutoff: 10

After alignment, a mean of 1.68 and 1.56 million uniquely mapped reads was achieved in the cross-sectional cohort and diurnal cohort, respectively (QC metrics in Supplementary Table 7).

### Sample exclusion

Of the 150 individuals recruited in year 1 of the cross-sectional cohort, 5 were excluded for lack of a repeat sampling in year 2. Sequencing libraries underwent quality control assessment prior to inclusion. Quality metrics included enzymatic conversion efficiency (>98%), evaluated by non-CpG methylation rates; detection of potential sample swaps through sex chromosome coverage consistency (chrX percentage in females and chrY percentage in males); and sequencing yield. One sample failed to meet these quality criteria and was excluded. This resulted in total of 288 samples from 144 individuals. All 144 diurnal samples met quality control criteria.

### cfDNA concentration estimation

Proxies for total cfDNA concentrations, expressed as spike-in to sample ratio, were generated based on read counts of lambda viral vector DNA, which was spiked into the library prior to processing. Concentration estimates were adjusted for duplication rate. Genomic equivalence was calculated using the following formula:

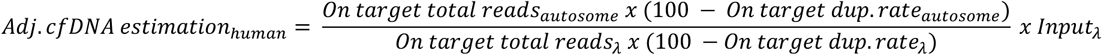

Total cfDNA concentrations were measured with Qubit (1X dsDNA HS Assay Kit, Thermo Fisher Scientific) post-extraction according to manufacturer’s guidelines for a subset of samples (*n =* 23). These showed adequate correlation (R = 0.83*, P* < 0.001) with lambda read-based estimated concentrations (Supplementary Figure 1a).

### Methylation-based cell type deconvolution

Reference DNA methylation data from GSE186458^12^ and GSE206818^63^ were used for cell-type-specific marker selection. Reference samples with low purities were excluded. A total of 103 samples from 24 cell types were selected. Sex chromosomes were excluded from the analysis. A one-versus-others strategy was used to derive Benjamini-Hochberg adjusted *P* values from two-sample t-test and absolute methylation differences between two groups. Both *P* values and absolute differences were ranked and the top 300 markers for each cell type with either adjusted *P* values smaller than 0.05 or absolute differences greater than 0.2 were retained. Markers that were not unique to a specific cell type were further excluded, resulting in 5,699 markers for deconvolution analysis. EpiDISH was applied for cell type deconvolution^26^. Major contributing cell types were defined as granulocytes, monocytes, megakaryocytes, erythrocyte progenitors, NK cells, CD4+ T cells, B cells, hepatocytes and vascular endothelium. Given the strong positive correlation between megakaryocytes and erythrocyte progenitors, and their shared origin from the megakaryocyte-erythroid progenitor (MEP) lineage, we combined the two cell types in MEP for cell type-specific effects on fragmentation analyses.

### cfDNA methylation-based age prediction model

Methylation levels from each CpG in the capture set were correlated with age from the year 1 dataset using Spearman’s rank correlation. CpGs with an absolute correlation no less than 0.1 were selected for model training. 9,275 selected CpGs and sex were used as variables, where sex was coded as non-penalized factor. Using 144 samples from year 1, leave-one-out procedures were applied to train elastic net regression models and to tune parameters. A final of 146 CpGs were selected to build the final model together with sex. This final model was applied on samples from year 2 to assess the prediction.

### End motif analysis

The terminal 2-nucleotide sequences at the ends of DNA fragments were extracted based on alignments to the reference genome. Frequencies of all possible 2-mer were calculated across fragments from a sample. 2-mer end motif frequencies for highly or lowly methylated regions were calculated based on fragments that overlapped with the defined regions.

### Marker-based fragment assignments

To assign cfDNA fragment to specific cell types, we employed a marker-based strategy using cell-type-specific DNA methylation signatures. A subset of cell-type-specific CpGs was selected from the 5,699 deconvolution markers. For each candidate CpG, we first calculated the average methylation level per cell type. A site is classified as a cell-type-specific marker if it showed a minimum absolute methylation difference of 0.6 between the cell type of interest and all other cell types. Markers were categorized as hypomethylated markers (the cell type of interest exhibited methylation at least 0.6 lower than the minimum averaged methylation observed in other cell types) or hypermethylated markers (the cell type of interest exhibited methylation at least 0.6 higher than the maximum averaged methylation in other cell types). For fragment assignment, cfDNA fragments covering a given marker were considered cell-type-specific if their methylation state matched the expected methylation level – 0% methylation for hypomethylated markers or 100% methylation for hypermethylated markers.

Fragments spanning multiple markers with discordant methylation states were excluded to ensure assignment specificity. In comparing fragment size of a cell type of interest to others, fragment size distributions of each cell type were contrasted against an aggregated background composed of fragments assigned to all other cell types.

### In vitro size selection analysis

Fragments falling within predefined size ranges (<167, 167-250, or >250 bp) were extracted from the aligned BAM files based on insert size. Samples with less than 10-fold coverage after size selection were excluded from downstream analyses. Methylation extraction was then performed on the size-selected BAM files, and cell type deconvolution was performed using the same procedure and marker set as in the analysis without size selection. Cell type contributions from size-selected and unselected samples were compared in a paired manner to assess the effect of fragment size on deconvolution outcomes.

### Canonical correlation analysis

A regularized canonical correlation analysis to explore relationships between major cell type proportions and cfDNA 2-mer end motif frequencies was performed. Both matrices of cell type proportions and end motif frequencies were standardized. Regularization parameters were tuned via cross-validation. Permutation tests that shuffled the data matrix of cell type proportions for 1,000 times were performed to validate whether correlations are non-random. Loadings that identify which cell types/motifs contribute most to the first two dimensions were shown. Projections of the original data into the first two CCA-derived latent spaces were extracted to visualize the association between cell types and motifs.

### Statistical analysis

Coefficient of variations were calculated as the ratio of the standard deviation to the mean. Repeated measures ANOVA tests, using a within-subject design, were applied to examine the effects of time and day in concentration or fragment size analysis of the diurnal cohort. Linear mixed effects models were adopted to examine both fixed and random effects in repetitive measurements. The Shapiro-Wilk test was used to assess normality assumptions. Log10 transformation was applied to the response variable where the normality assumption was not met. Individuals, time of sampling and year of sampling were taken as random variables. The variance-covariance components of the random effects were calculated to derive intraclass correlation coefficients. Paired t-tests were used to compare the means of end motif frequencies between highly and lowly methylated regions and deconvolution results between size-selected and unselected paired samples. To assess correlations between different cfDNA fragmentation characteristics, Spearman’s rank correlation coefficients were reported. Kendall rank correlation coefficients were reported in assessing inter- and intra-individual cell type and fragment size variations. Pearson’s correlation coefficients were reported in size-size and motif-motif correlations.

## DECLARATIONS

### ETHICS APPROVAL

The study (no. S66450) was approved by the Medical Ethics Committee of University Hospitals Leuven.

### DATA AVAILABILITY

The datasets generated and/or analysed during the current study will be available in the European Genome-Phenome Archive (EGA) repository under controlled access.

### COMPETING INTERESTS

B.T. received Illumina speaking fees for work unrelated to the work presented here.

## FUNDING

The resources and services used in this work were provided by the VSC (Flemish Supercomputer Center), funded by the Research Foundation - Flanders (FWO) and the Flemish Government. This study was supported by FWO-SBO grant S003422N, FWO grant G0B4822N to B.T., and institutional support from KU Leuven, C1- C14/22/125 to J.R.V, METH/21/06 to B.T., and C3 - C3/24/078 to B.T. and J.R.V.. M.A. is supported by an FWO fellowship.

## AUTHORS’ CONTRIBUTIONS

B.T., J.R.V., T.J., K.D. and M.A. conceptualized and designed the study. B.T., M.A., T.J. and J.R.V. organized and supervised the project. M.A., T.J., K.L., A.N. and V.P. were involved in the sample processing pre-sequencing. K.D.R. developed the capture set and K.D.R. and H.C. performed post-sequencing quality control. H.C. performed all bioinformatics and statistical analyses. B.T., H.C., and M.A. wrote the manuscript. All coauthors reviewed the manuscript.

## ACKNOWLEDGEMENTS

We would like to thank Marian Crabbé for her support in preparing and submitting the ethics application, Albert Herelixka, Angelica Pagliazzi, Stefania Tuveri and Miel Theunis for their assistance in coordinating study visits and sampling study participants, and Leen Vancoillie and the prenatal diagnostics team at the Leuven University Hospital for cfDNA extraction. Finally, we thank all study participants for their interest and enthusiasm.

## FIGURE LEGENDS

**Supplementary Figure 1:**
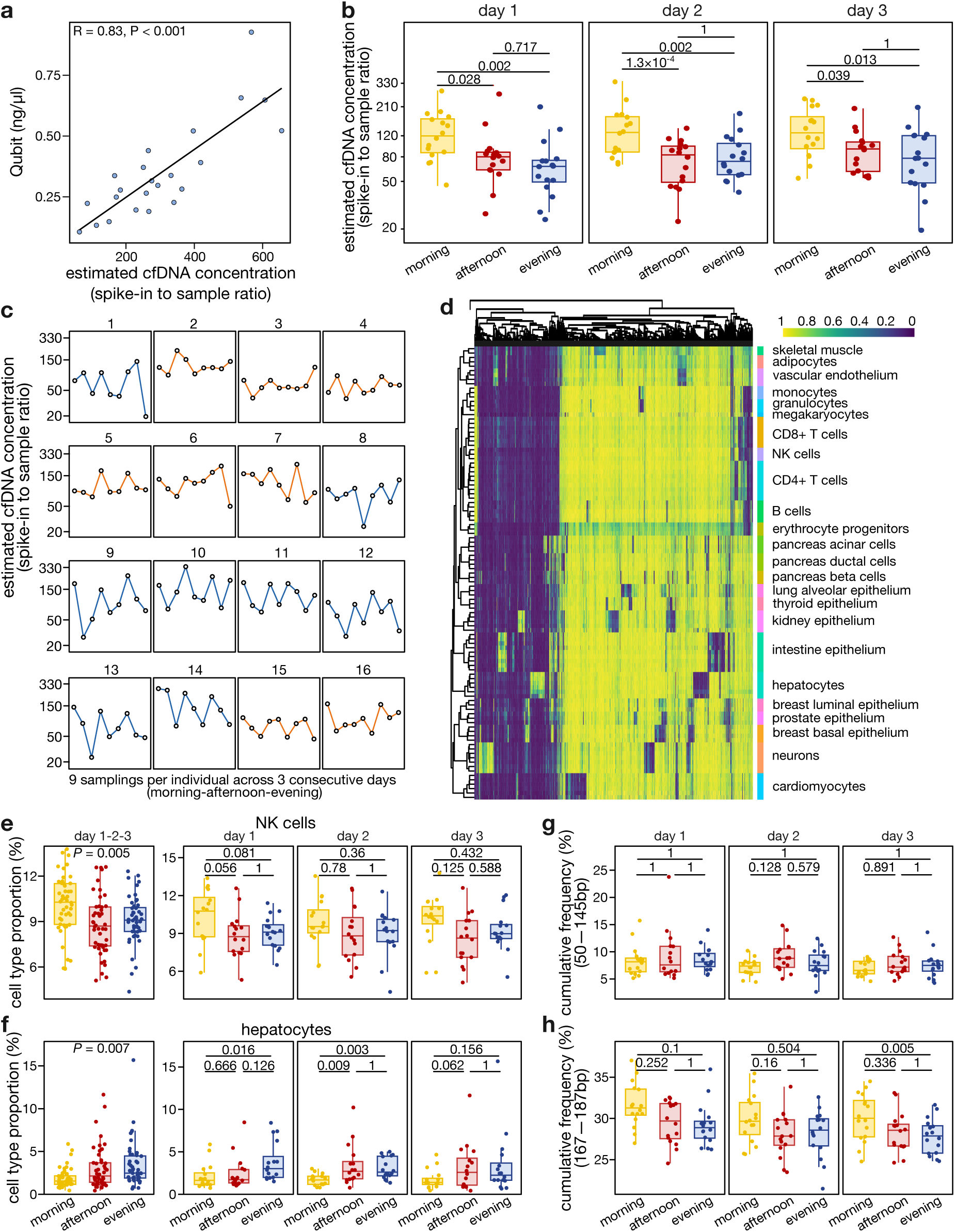
Diurnal variation in cfDNA characteristics. **a,** Pearson correlation between cfDNA concentrations measured by Qubit post-extraction (*n* = 23 samples) and estimated concentrations based on spiked-in lambda viral vector DNA (Methods). **b**, Estimated cfDNA concentration levels across timepoints (morning, afternoon and evening) over three consecutive sample days. **c**, Individual estimated cfDNA concentration trajectories for male (blue) and female (orange) participants in the diurnal cohort across the three sampling days. **d**, Heatmap of marker CpG methylation (*n* = 5,699) used for cell type deconvolution of 24 cell types. **e-f,** Diurnal variation in cell type proportions for NK cells **(e)** and hepatocytes **(f)** in samples combined per timepoint (left) and across the three sampling days (right). **g-h**, Cumulative frequency of subnucleosomal (**g**; 50-145 bp) and mononucleosomal (**h**; 167-187 bp)-sized fragments at different timepoints over three consecutive sampling days. *P* values by Pearson correlation (a), repeated measures ANOVA for all sampling days combined, followed by pairwise t-test on each individual day (b, e, f, g, h).

**Supplementary Figure 2:**
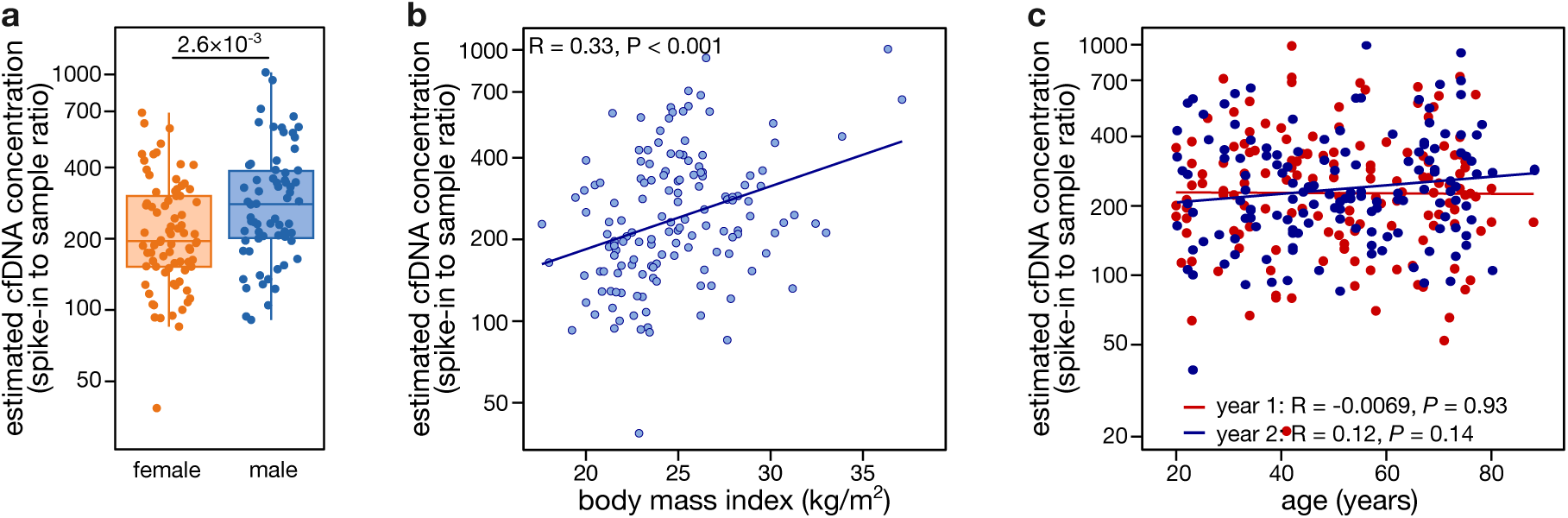
Demographic factors affecting cfDNA concentration. **a**, Comparison of estimated cfDNA concentrations (log10 scaled) between female and male participants in year 2. **b**, Pearson correlation between body mass index and estimated cfDNA concentration (log10 scaled) in year 2. **c**, Pearson correlation between age and estimated cfDNA concentration (log10 scaled) in year 1 (red) and year 2 (blue). *P* value by Wilcoxon rank-sum test (a) or Pearson correlation (b, c).

**Supplementary Figure 3:**
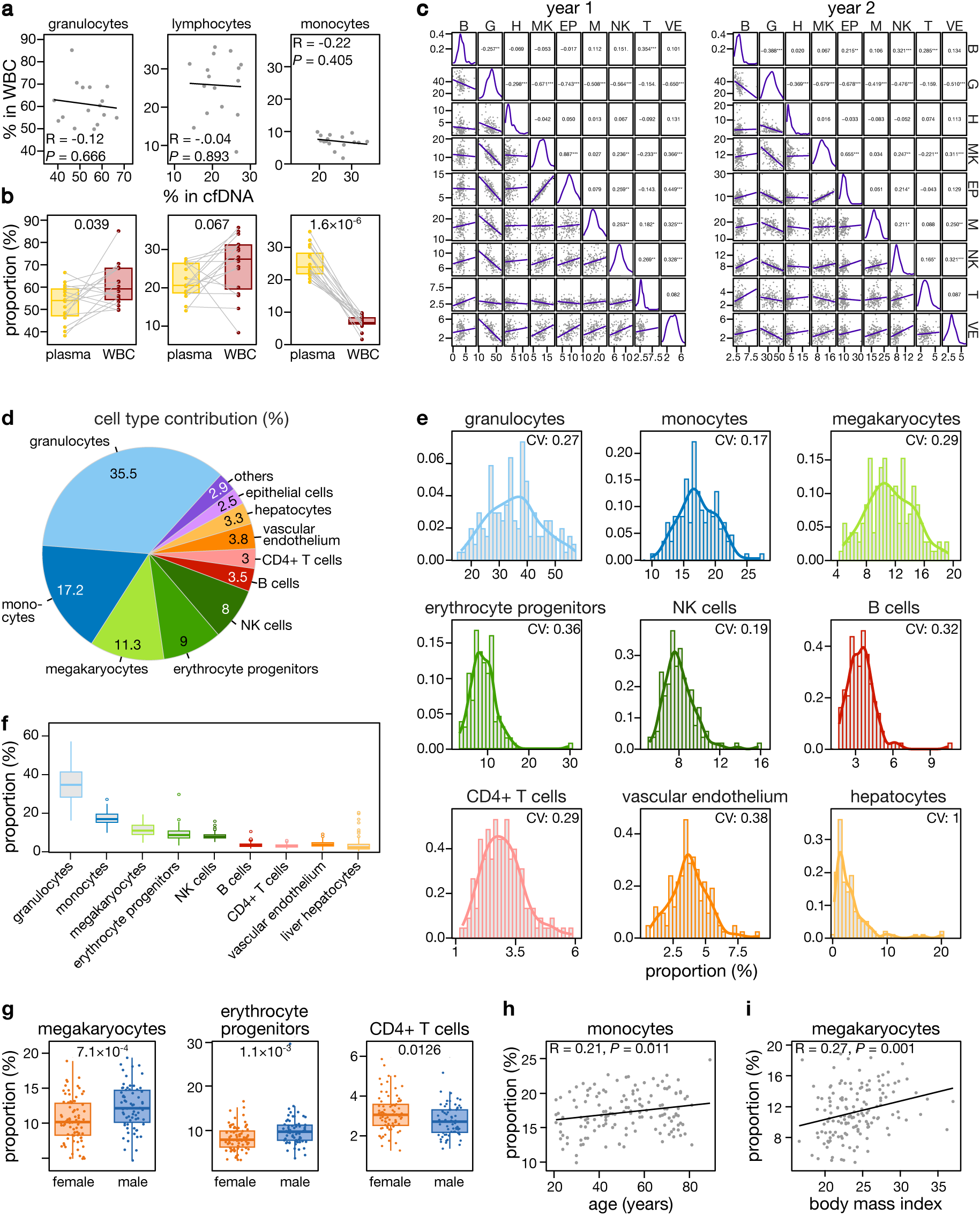
Factors influencing plasma cfDNA composition. **a**, Pearson correlation between cfDNA-derived white blood cell (WBC) proportions and corresponding cell type proportions in WBCs from whole blood. In whole blood, neutrophil proportions are used as a proxy for granulocytes, given the typically low levels of eosinophils and basophils. **b**, Comparison of cfDNA-derived WBC proportions with matched proportions in WBCs from whole blood. **c**, Pearson correlation between cfDNA fractions for different contributing cell types in year 1 and 2. Significance codes: <0.001 ‘***’, <0.01 ‘**’, <0.01 ‘*’, <0.05 ‘.’, 1 ‘’. **d**, Plasma cell type contributions to cfDNA in year 2 samples, represented as a pie chart. Other cell types include adipocytes, cardiomyocytes, neurons, pancreatic cells, and skeletal muscle cells. **e**, Distribution of proportions (density) for major cell types contributing to plasma cfDNA in year 2. **f**, Interindividual variation in major cell types contributing to plasma cfDNA in year 2. **g**, Comparison of cell type contributions to plasma cfDNA between female and male participants in year 2. **h**, Pearson correlation between monocyte proportions and age in year 2 samples. **i**, Pearson correlation between megakaryocyte proportions and body mass index (kg/m^2^) year 2 samples. B: B cells; G: granulocytes; H: hepatocytes; MK: megakaryocytes; EP: erythrocyte progenitors, M: monocytes; NK: NK cells; T: CD4+ T cells; VE: vascular endothelium. *P* values by Pearson correlation (a, h, i), paired t-test (b), Pearson correlation (c) or Wilcoxon rank-sum test (g).

**Supplementary Figure 4:**
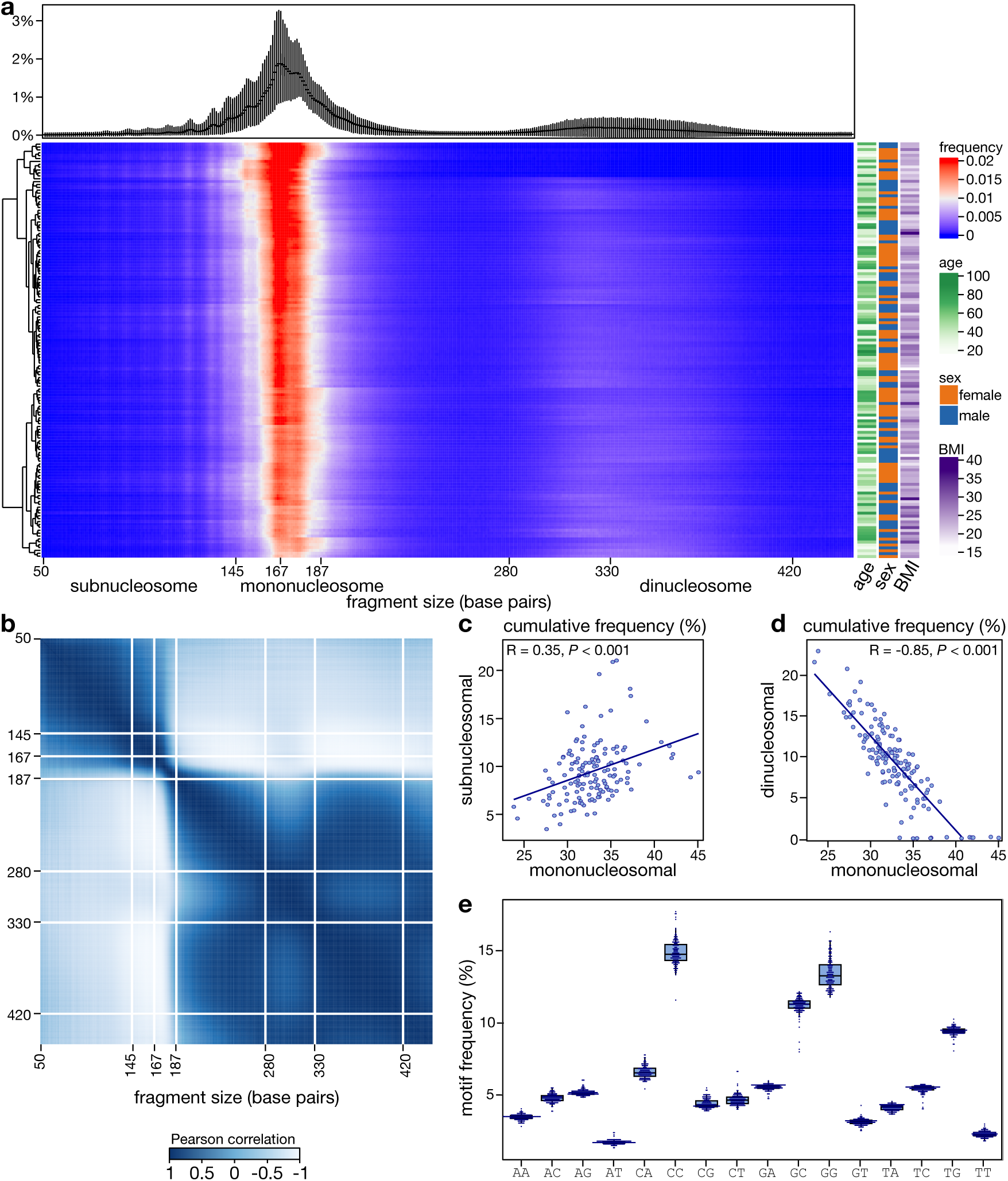
cfDNA fragmentation patterns in the cross-sectional cohort. **a**, cfDNA fragment size distributions in year 2 samples, shown as average profiles with error bars representing upper and lower quantiles (top), and a correlation matrix showing associations between fragment sizes and demographic variables (sex, age, and BMI; bottom right). Fragment sizes are defined as subnucleosomal (<145 bp), mononucleosomal (167-187 bp) and dinucleosomal (330-420 bp). **b**, Overview of fragment size correlations in year 2 samples, demonstrating a positive correlation between mononucleosomal and subnucleosomal fragments (**c**), and a negative correlation between mononucleosomal and dinucleosomal fragments (**d**). **e**, Relative contributions of dinucleotide end motifs to plasma cfDNA in year 2 samples. *P* values by Pearson correlation (c, d).

**Supplementary Figure 5:**
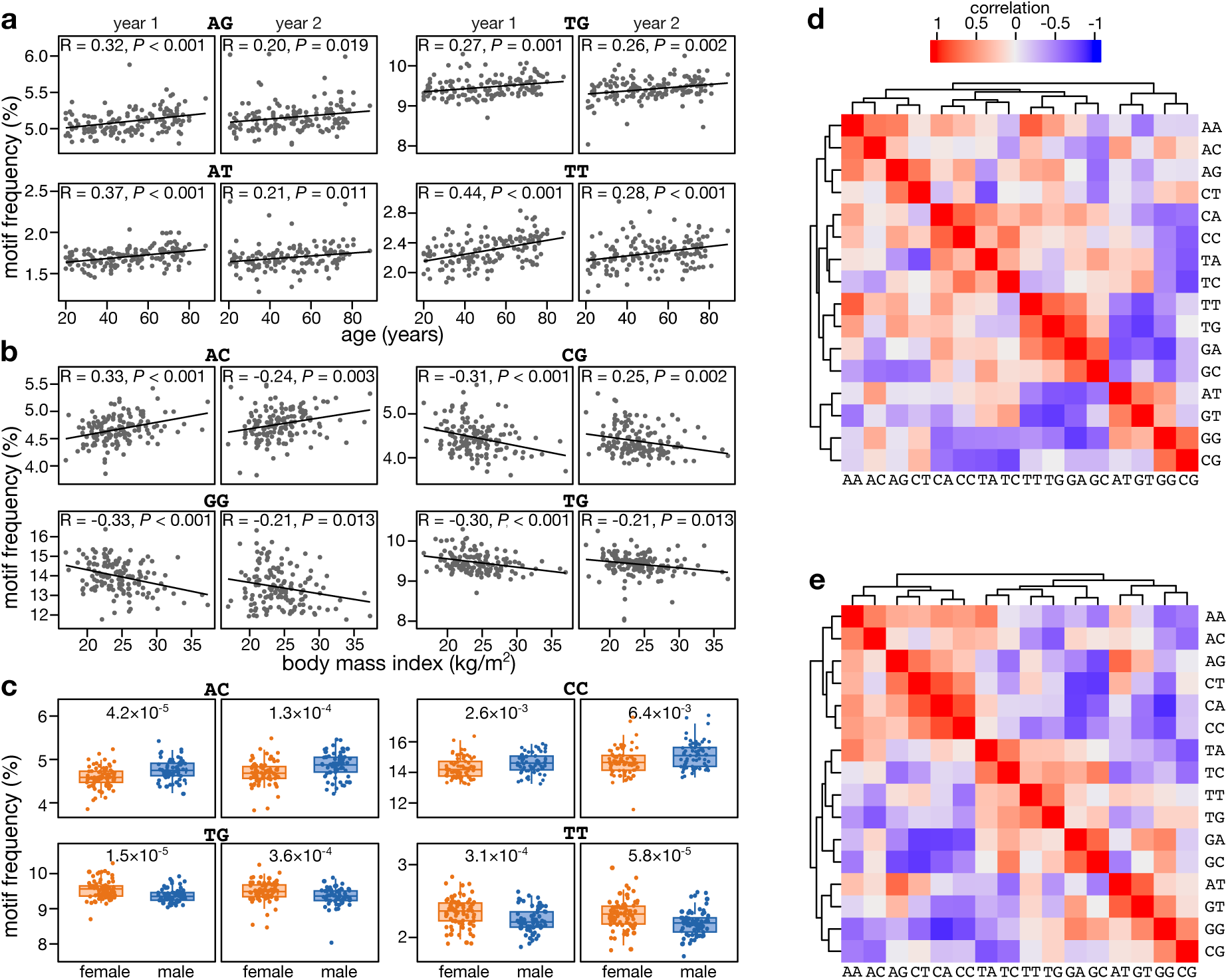
**a-c**, Associations between dinucleotide end motif frequencies and demographic variables in year 1 and year 2 samples. Shown are correlations with age (a), BMI (b) and sex (c). **d**,**e**, Overview of dinucleotide end motif correlations (Pearson) in year 1 (d) and year 2 (e) samples. *P* values by Pearson correlation (a, b), Wilcoxon rank-sum test (c).

**Supplementary Figure 6:**
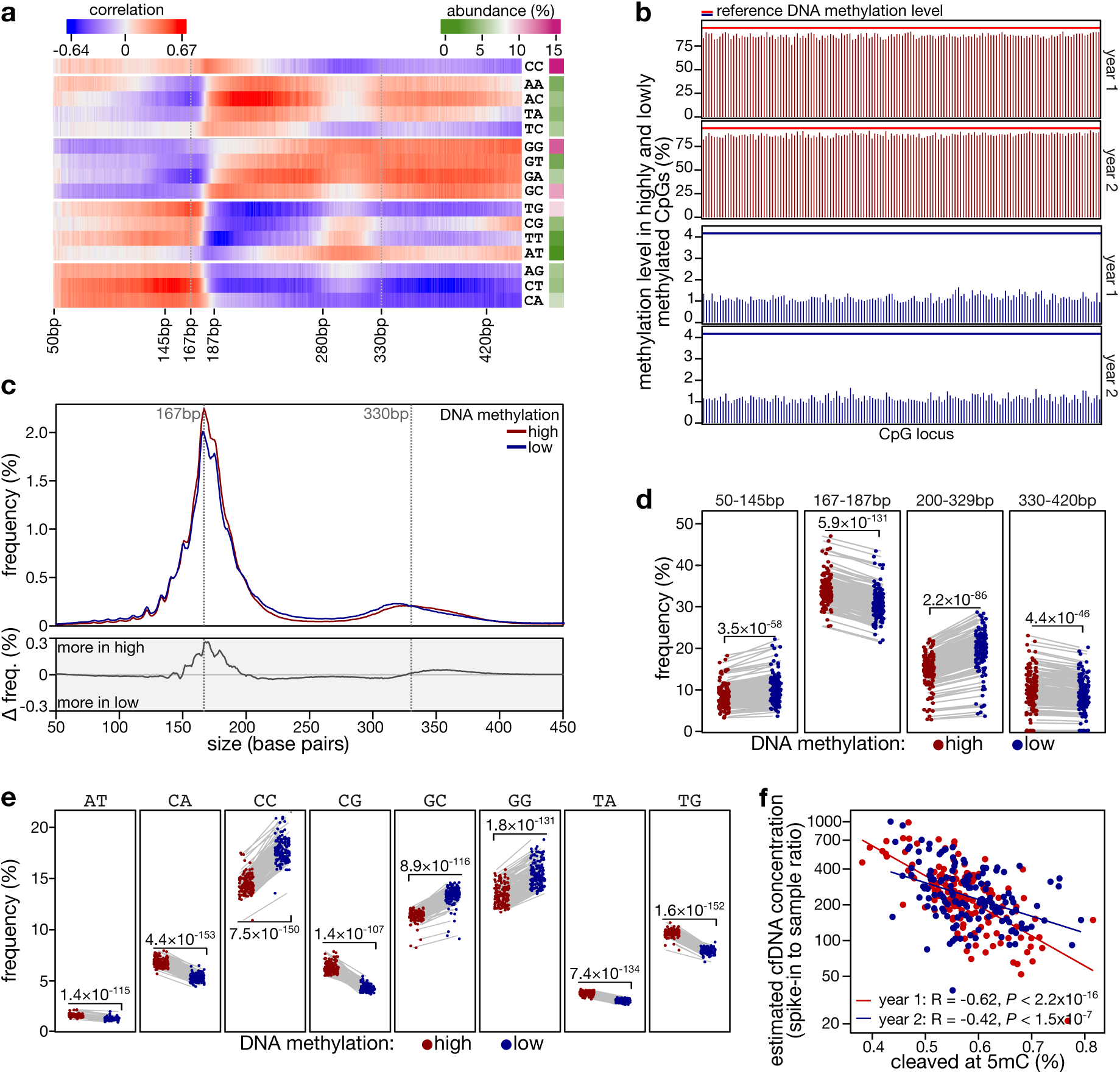
Factors affecting cfDNA fragmentome characteristics. **a**, Spearman correlation matrix showing associations between cfDNA fragment sizes and 2-mer end motifs in year 2 samples, for subnucleosomal (<145 bp), mononucleosomal (167-187 bp), and dinucleosomal fragments (330-420 bp). **b**, CpG loci classified as highly methylated (top, *n* = 2,953) or lowly methylated (bottom, *n* = 2,752) across major contributing cell types (see Methods). Reference methylation indicates expected levels at each locus; bars show corresponding methylation levels in cfDNA. **c**, Fragment size distributions (top) and frequency difference (bottom) of cfDNA fragments overlapping the highly and lowly methylated CpGs defined in (b) in year 2 samples. **d**, Cumulative frequency of highly and lowly methylated fragments across different fragment size ranges in year 2 samples. **e**, Frequency of highly and lowly methylated fragments across different 2-mer end motifs in year 2 samples. **f**, Pearson correlation between cleavage at cytosine in CpG context for highly methylated fragments and estimated cfDNA concentration. bp: base pairs. *P* values by paired t-test (d, e) and Pearson correlation (f).

**Supplementary Figure 7:**
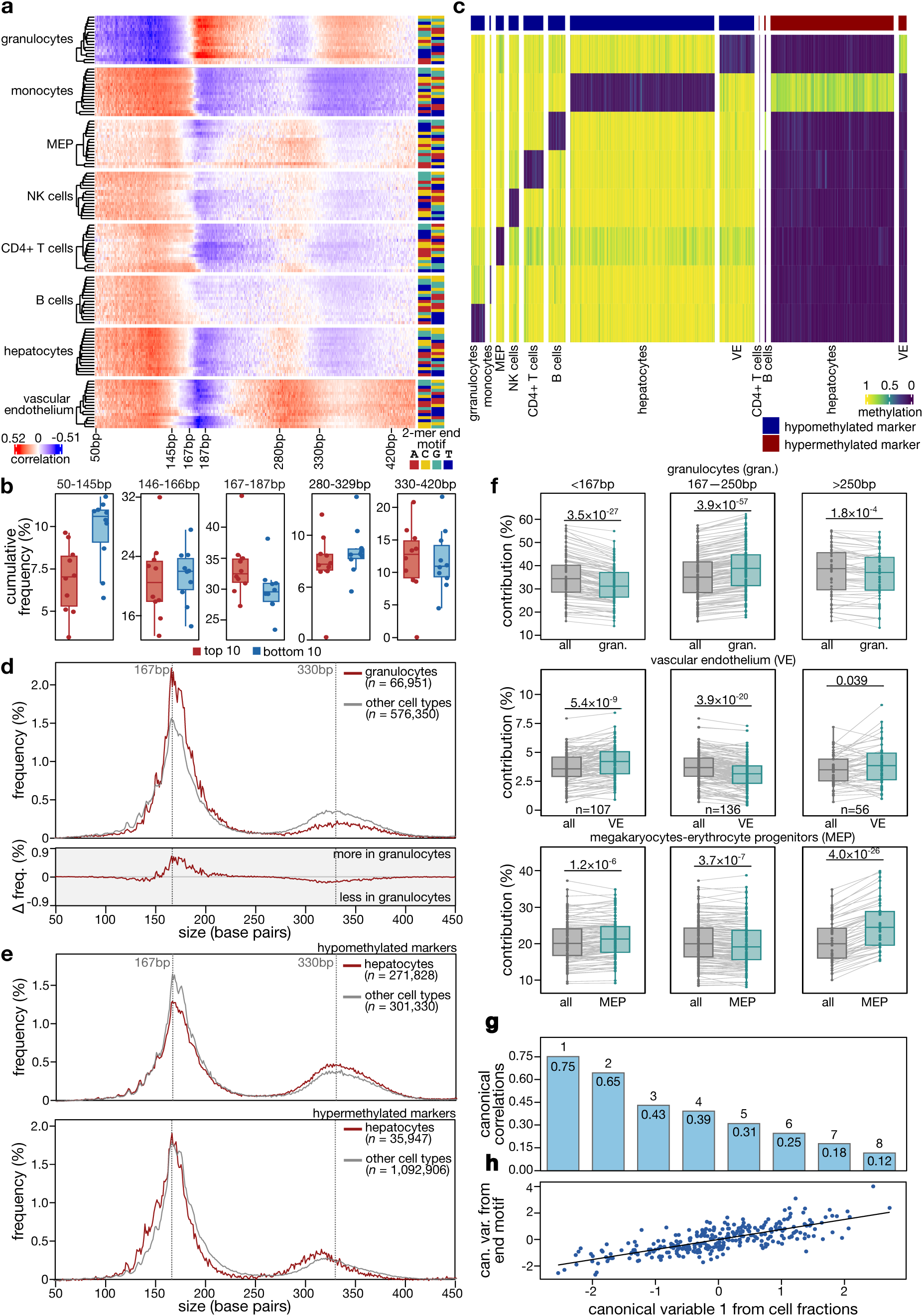
Cell-type-specific effects on cfDNA fragmentation. **a**, Spearman correlation matrix showing cell-type specific associations between cfDNA fragment sizes and 2-mer end motifs for major contributing cell types in year 2 samples. **b**, Cumulative frequency of fragments across different fragment size ranges in the top ten samples with the highest versus lowest granulocyte proportions in year 2. **c**, Heatmap of cell type-specific DNA methylation markers showing at least a 60% difference in average methylation relative to other cell types. **d**, Fragment size distributions (top) and frequency difference (bottom) between hypomethylated granulocyte-specific fragments and all other hypomethylated cell type-specific fragments in year 2 samples. **e**, Fragment size distributions of hypo- and hypermethylated hepatocyte-specific cfDNA fragments (red) compared to those of other cell types (grey) in year 1 samples. **f**, Deconvolution analysis of cfDNA from year 2 samples after *in silico* size selection into three bins (<167 bp, 167–250 bp, >250 bp); samples with <10× coverage per bin were excluded, retaining 107, 136, and 56 samples per bin respectively. **g**, Canonical correlations showing in all canonical dimensions. **h**, Correlation between cfDNA fragmentation patterns and cell type composition along canonical variable 1. *P* values by paired t-test (f).

